# Lower spatial turnover of rare fungal taxa dominantly shaped by stochastic processes in grassland soils

**DOI:** 10.1101/2025.01.20.633927

**Authors:** Zhang Biao, Zhou Shutong, Xue Kai, Liu Wenjing, Chen Shuo, Tang Li, Li Linfeng, Du Jianqing, Hao Yanbin, Cui Xiaoyong, Wang Yanfen

**Affiliations:** College of Resources and Environment, University of Chinese Academy of Sciences, Beijing 100049, China; State Key Laboratory of Plateau Ecology and Agriculture, Qinghai University, Xining 810016, China; Beijing Yanshan Earth Critical Zone National Research Station, University of Chinese Academy of Sciences, Beijing 101408, China; College of Life Sciences, University of Chinese Academy of Sciences, Beijing 101408, China; State Key Laboratory of Tibetan Plateau Earth System Science (LATPES), Beijing 100101, China

**Keywords:** Abundant fungal taxa, Rare fungal taxa, Distance decay, Spatial turnover, Community assembly

## Abstract

The spatial pattern and community assembly of soil microbial taxa have notable meanings for biodiversity shaping and maintaining mechanisms. Rare fungal taxa may exhibit distinct patterns and assembly mechanisms compared to abundant taxa, but such information is limited, especially at large scales. Here, we investigated distance-decay patterns and underlying assembly mechanisms for abundant and rare fungal taxa in 129 soil samples collected across 4,000 km in Chinese Northern grasslands, based on high-throughput sequencing data. A total of 208 abundant OTUs (relative abundance > 0.1%, 2.73% of entire OTUs) and 5,779 rare OTUs (relative abundance < 0.01%, 75.85% of entire OTUs) were identified. Both abundant and rare fungal taxa showed significant distance-decay relationships (P < 0.001), but the turnover rate for rare taxa (0.0024 per 100 km) was nearly half that of abundant taxa (0.0054 per 100 km) based on the binary Bray-Curtis distance. The lower turnover of rare fungal taxa was likely due to their community assembly mechanism dominated by stochastic processes, which were less influenced by environmental gradients. In contrast, abundant taxa assembly was dominated by deterministic factors like soil variables and plant traits, which varied significantly along the geographic distance. Consistently, rare fungal taxa were also less sensitive to environmental changes, with a lower turnover rate by environmental distance (0.0027 vs. 0.0099) than abundant taxa. In summary, our findings revealed that rare fungal taxa, shaped mainly by stochastic processes, had lower spatial turnover compared to abundant taxa, dominated by deterministic processes, enhancing our understanding of rare microbial biogeography.

## Introduction

The spatial pattern of soil microbial community has received increasing attention since the advent of molecular biology tools and high-throughput sequencing technology, which empowered researchers to capture the microbial diversity than ever (Caporaso et al. 2010). However, most of previous microbial geographical studies focused on bacteria or prokaryotes (Bell 2010, Locey et al. 2020, Zhang et al. 2022), but ignored fungi likely due to the complexity of their classification. As one of the most abundant and diverse biological groups on the earth (Pedros-Alio 2012, Chen et al. 2020), soil fungi play fundamental roles in driving soil nutrients cycling, mediating plant mineral nutrition, and alleviating nutrient limitations (Hug et al. 2016, Jiao and Lu 2020a), or leading to plant disease (Bever 2015). Fungi differ greatly from bacteria in terms of individual morphology, mobility, and substrate utilization (Tedersoo et al. 2014, Peay et al. 2016) etc. For example, the special hyphal filaments of some fungi facilitate it capturing more resources from heterogeneous environments at the micrometer scale (Jiao and Lu 2020a) than bacteria or prokaryotes do. These different features of fungi may lead to a distinct spatial pattern and community assembly from that of bacteria or prokaryotes. Conflicting results were obtained from a few existing studies, which have shown that soil fungal community exhibits distance-decay patterns with higher spatial turnover rates than those of the bacterial community at scales from hundreds of meters to thousands of kilometers in agriculture ecosystems (Jiao and Lu 2020a, Ma et al. 2023b) and lake sediments (Wang et al. 2023b), but with lower turnover rates at a scale over 3, 000 km in boreal forest soils (Ma et al. 2017).

Microbial community members could be classified into abundant and rare species according to individual species’ abundances, judged by the threshold of 0.1% or 0.01% (Jia et al. 2018, Pascoal et al. 2021). Microbial communities harbor a large number of rare taxa coexisting with a small fraction of abundant ones, though a small proportion of rare taxa periodically or occasionally shifted to high abundances in response to environmental changes (Shade et al. 2014, Pascoal et al. 2021). As the primary component of microbial communities, abundant taxa usually distribute broadly and exhibit certain tolerance to environmental changes (Delgado-Baquerizo et al. 2018).

Though most of our microbial knowledge is based on abundant species, increasing attention has been paid on the microbial rare biosphere recently (Jousset et al. 2017, Tang et al. 2023, Wang et al. 2024). Despite their low abundances, rare taxa may perform over-proportional roles in maintaining soil biogeochemical cycles, biodiversity and ecosystem stability (Chen et al. 2020), working as a hidden driver of microbial functions (Jousset et al. 2017). Rare taxa has been reported to hinder the establishment of an alien species through resource competition (Mallon et al. 2015). They serve as reservoirs of species (Jia et al. 2018) and contribute greatly to the quantification of species richness in a given community. Based on bacterial communities, the high richness and low abundance of rare taxa had different spatial distributions from abundant taxa or the entire community in diverse environments like contaminated soils (Jiao et al. 2017), subtropical bays (Mo et al. 2018), agriculture ecosystems (Ma et al. 2023a) and desert soils (Sun et al. 2023), and were identified as a driver for microbial variation in space and time (Alonso-Saez et al. 2015, Chen et al. 2020, Wang et al. 2023a). Conflicting results were observed in previous reports, as some found that the distance-decay patterns for rare taxa exhibited insignificant or weak relationships compared to those of abundant taxa in agriculture fields (Jiao and Lu 2020a), deserts (Wang et al. 2022a) and acidic soils (Wang et al. 2023a), while limited explorations also revealed higher turnover rates for rare than abundant fungal taxa (Jiao and Lu 2020a, Wang et al. 2022a, Tang et al. 2023, Wang et al. 2023a). So far, there is still a lack of knowledge on the distance-decay relationship and its turnover rate of rare fungal taxa, and what determines such spatial pattern.

Understanding underlying mechanisms controlling the microbial spatial pattern is a central but poorly understood topic in microbial ecology. Roles of stochastic and deterministic processes in shaping microbial spatial distribution are increasingly recognized (Zhou and Ning 2017, Jia et al. 2018, Wang et al. 2023c). From the aspect of deterministic processes, which are represented by the niche theory (Chesson 2000, Fargione et al. 2003), environmental conditions and species interactions could determine the community diversity and spatial patterns. The stochastic processes, which are represented by the neutral theory, are based on the “ecological equivalence” and believe that stochastic processes such as birth, death, migration, extinction, drift, and speciation determine the community diversity (Stegen et al. 2013, Zhou and Ning 2017). Some empirical evidence suggests that deterministic and stochastic processes work simultaneously to shape the community structures but with different contributions, likely depending on ecosystem types or geographic scales (Shi et al. 2018, Pascoal et al. 2021, Tang et al. 2023). Stochastic processes were considered to determine predominantly fungal communities’ distribution patterns at the local or regional scales (Yang et al. 2017). Deterministic processes, including soil pH (Zhang et al. 2018, Wang et al. 2023a), water (Li et al. 2017) and nutrients (Zhao et al. 2022b), have been found to be more important in driving soil fungal communities in scales from hundreds to thousands kilometers. Conflicting results have been found in previous studies for the community assembly of rare and abundant taxa, dominated by similar (Wan et al. 2021, Zhao et al. 2022a, Tang et al. 2023) or disparate processes (Xue et al. 2018, Ji et al. 2020, Jiao and Lu 2020a). For example, a study in the Tibetan alpine grassland found that deterministic processes dominate either rare or abundant fungal taxa, but the composition of rare rather than abundant fungal taxa associated with the plant community (Tang et al. 2023).

Information regarding the distribution pattern and underlying mechanisms of abundant and rare fungi taxa at a large scale is still lacking in grassland soils, which are essential for understanding biodiversity shaping and maintaining mechanisms (Jiao and Lu 2020a). The present study explored the distance-decay relationships of the entire, abundant and rare fungal taxa and their community assembly mechanisms by surveying soil samples from a transect across 4,000 km in the Chinese northern grassland. We hypothesized that (1) both abundant and rare fungal taxa would exhibit distance-decay patterns, but with different turnover rates, and (2) the community assembly of abundant and rare fungal taxa would be determined by different processes.

## 2 Materials and Methods

### 2.1 Data collection

Soil samples were collected from July 29 to August 14, 2014 in the Qinghai-Tibet Plateau alpine grassland (a.s.l. 2,796-4,891 m) and from September 10 to 24, 2015 in the Inner Mongolia temperate grassland (a.s.l. 10-1,796 m) of China as described by Zhang et al. (2022). A total of 128 samples from 64 sites were collected in the alpine grassland and 130 samples from 65 sites were collected in the temperate grassland. The distance between each two adjacent sample sites was more than 60 km. Aboveground and belowground (0-5 cm in depth) plant parts were collected and dried at 65°C for 72 h as aboveground biomass (AGB) and belowground biomass (BGB), respectively. Soil pH, water content (SWC), soil organic carbon (SOC), dissolved organic carbon (DOC), total nitrogen (TN), dissolved organic nitrogen (DON), total phosphorus (TP), available phosphorus (AP), ammonium (NH_4_^+^), nitrate (NO_3_^-^) were measured and the climate data of mean annual temperature (MAT) and mean annual precipitation (MAP) were collected by latitude and longitude from ‘China Meteorological Data Service Center’ (CMDC: https://data.cma.cn/). Environmental variables were divided into long-term environmental variables and short-term environmental variables by whether they remain stable within 1 year or not (Zhang et al. 2022).

Soil fungal diversity was analyzed using high-throughput sequencing of the internal transcribed spacer (ITS2) region with the primer pair ITS4F (5’-TCCTCCGCTTATTGATATGC-3’) and gITS7R (5’-GTGARTCATCGARTCTTTG-3’) on a MiSeq Platform (Illumina Inc.). The MiSeq raw data was analyzed by UPARSE pipeline with USEARCH 8 (Edgar 2013) software to obtain an operational taxon units (OTU) table at the threshold of 97% identity. Each OTU was annotated by Mothur (v1.27) (Schloss et al. 2009) with classify.seqs command, and UNITE was selected as the reference database. The OTU table was resampled to the same sequence (8,323) before further analysis by R 4.3.0 with the resample package. The OTUs with relative abundances > 0.1% of the total sequences were defined as “abundant” OTUs (Jiao and Lu 2020a), and all abundant OTUs composed to abundant taxa. The OTUs with relative abundances < 0.01% were defined as “rare” OTUs (Jia et al. 2018, Pascoal et al. 2021), and 0.1% were “intermediate” OTUs.

### 2.2 Data analysis

Non-metric multidimensional scaling (NMDS) analysis based on Bray-Curtis distance and Venn analysis were performed to test the effect of biotas on the compositions of soil entire fungal community, abundant fungal taxa and rare fungal taxa. The geographic distance (km) between each two sites was calculated based on their longitude and latitude using the “distGeo” function of “geosphere” package in R program, which considers spheric deviations (Zhang et al., 2022). The similarity and dissimilarity of plant community were calculated based on Bray-Curtis distance by the vegan package of R. The similarity and dissimilarity of the entire, abundant and rare fungal taxa were calculated based on binary Bray-Curtis distance to eliminate the effect of abundance that differed greatly between them inevitably. All environmental variables were normalized before calculating the Euclidean distance. Linear model and LSD were used to test the decay relationship of the fungal communities’ similarity, including entire, abundant and rare fungal taxa, based on binary Bray-Curtis distance with environmental Euclidean distance. Pearson correlation was used to test the relationship of soil fungal diversity with environmental variables. The Mantel test and Partial Mantel test based on a Pearson correlation were used to test the relationship of soil fungal similarity, geographic distance, environmental variables, and plant community distance. The turnover rate was estimated by the slope of the linear regression model based on the least square method.

The niche breadth of each OTU was calculated based on the Levins index, and the community niche (*B*) at each site was calculated by the niche of each OTU. According to the calculation formula of community niche breadth, the more dispersed the community distribution the greater its *B* value, and the more concentrated the community distribution the smaller its B value. Normalized stochastic ratio analysis (NST) was performed to distinguish the contribution of stochastic processes (homogenous dispersal, dispersal limitation, drift) and deterministic processes (homogenous selection and heterogeneous selection) to soil fungi community assembly.

The estimation of the immigration rate (*m*) based on Hubbell’s neutral theory of biodiversity was calculated by TeTame 2.0. Parameter estimation was rigorously performed by maximum-likelihood using the sampling formula developed by Etienne (50-53). In this model, the species relative abundances in a guild are determined by two parameters *θ* and *m*. The *θ* governs the appearance of a new species in the regional species pool, and *m* governs immigration into local communities of individuals from the regional species pool. Structural equation modeling (SEM) was performed by AMOS software to disentangle the causal pathways through which geographic distance, climate distance, edaphic distance, and plants’ community dissimilarity influence soil prokaryote similarity.

## 3 Results

### 3.1 Soil fungal diversity and community composition

A total of 24 phyla were detected (Figure S1), including *Ascomycota* (64.31 ± 19.53 % in relative abundance), *Basidiomycota* (15.97 ± 17.63 %), *Mortierellomycota* (14.85 ± 14.14 %), *Glomeromycota* (1.56 ± 2.2 %), *Chlorophyta* (1.21 ± 2.04 %), *Anthophyta* (0.53 ± 3.21 %). A total of 208 OTUs were classified as abundant fungal taxa with an average of 62.59 ± 15.02% sequences, accounting for 3.02% of total OTUs (6,886). In contrast, rare fungal taxa contained 5,779 OTUs with 8.75 ± 4.81% sequences on average, accounting for 83.92% of total OTUs. The Shannon diversities were 3.67 ± 0.83, 2.48 ± 0.66 and 4.18 ± 0.64 in all sites for entire, abundant and rare fungal taxa, respectively. Fungal communities’ Shannon diversities were lower in the alpine grassland (3.41± 0.85, 2.25 ± 0.67 and 4.05 ± 0.71 or entire, abundant and rare fungal taxa, respectively) than those in temperate grassland (3.93 ± 0.73, 2.70 ± 0.57 and 4.31 ± 0.54 for entire, abundant and rare taxa, respectively). The NMDS analysis showed that the compositions of entire fungal communities between alpine and temperate grasslands differed clearly (Figure S2a), and so did abundant (Figure S2b) or rare (Figure S2c) fungal taxa.

For entire fungal communities, according to the Venn plot, a total of 2, 760 OTUs were shared between alpine and temperate grasslands (Figure S2d), while 2, 148 and 1, 979 specific OTUs in alpine grassland and temperate grassland, respectively. However, the number of shared abundant OTUs was 179 between alpine and temperate grasslands, accounting for 88.61% of the abundant taxa (Figure S2e), while 1, 896 shared rare OTUs accounting for 32.81% of the total rare OTUs (Figure S2f). There were 15 and 8 specific abundant OTUs and 2,034 and 1,849 specific rare OTUs in alpine and temperate grassland, respectively.

### 3.2 Distance decay relationship for soil fungal taxa

On the scale of 4,000 km across alpine and temperate grassland biotas, the similarity of soil entire, abundant and rare fungal taxa decayed significantly along the increasing geographic distance (Figure 1), with rates of -0.0036 (R^2^ = 0.127, *P* < 0.001), -0.0051 (R^2^ = 0.094, *P* < 0.001) and -0.0021 (R^2^ = 0.120, *P* < 0.001) per 100 km, respectively.

**Figure 1.**
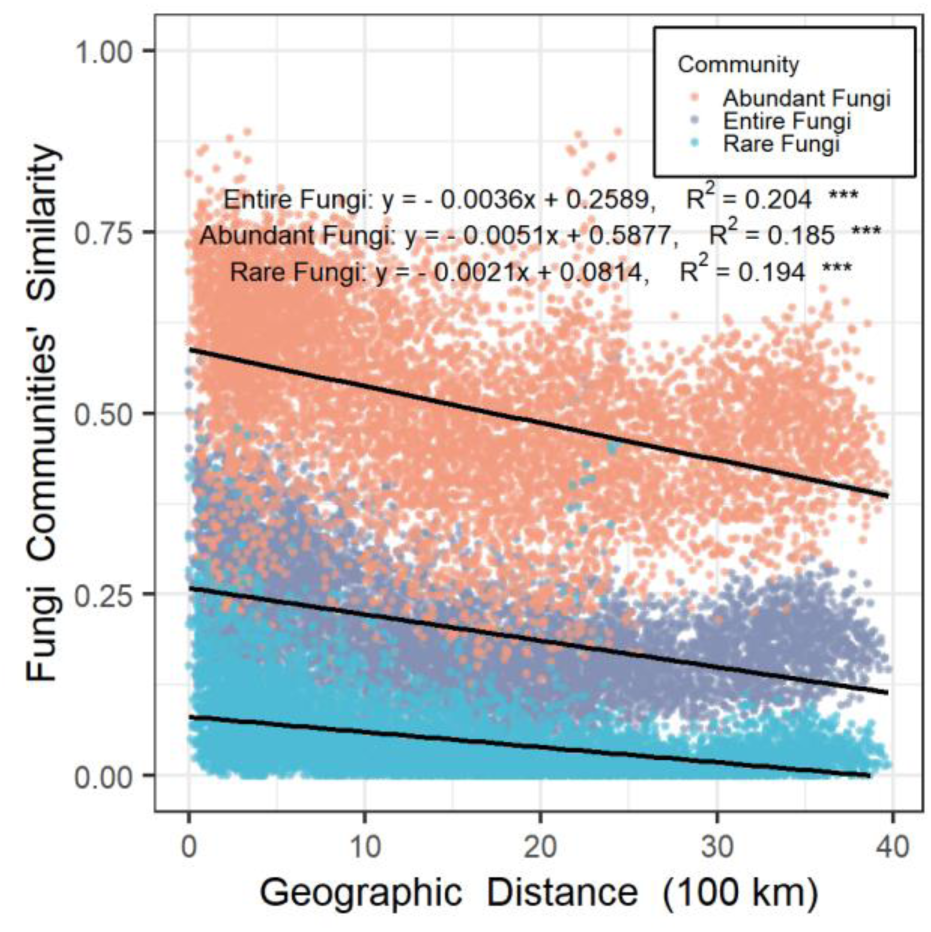
Similarities of soil entire, abundant and rare fungal taxa decreased with the increasing geographic distance across 4,000 km in both alpine and temperate grasslands. The grey, orange and blue points stand for entire, abundant and rare fungal taxa, respectively. “***” stands for significance at 0.001 level. Each fungal communities’ similarity was calculated based on binary Bray-Curtis index.

Similar decay patterns were observed in the alpine or temperate grassland alone, though the turnover rate for abundant fungal taxa was lower than that for entire and rare fungal taxa in the temperate grassland.

For the entire fungal taxa, the turnover rate was lower in the alpine grassland (-0.0058, R^2^ = 0.115, *P* < 0.001, Figure S3a) than that in the temperate grassland (-0.0065, R^2^ = 0.064, *P* < 0.001, Figure S3b). Similar pattern was also found for rare fungal taxa, though the turnover was higher in the alpine grassland (-0.0064, R^2^ = 0.049, *P* < 0.001, Figure S3a) than that in the temperate grassland (-0.0043, R^2^ = 0.039, *P* < 0.001, Figure S3b).

The dissimilarity of environmental variables increased with geographical distance (turnover rate = 0.0108 per 100 km, R^2^ = 0.0015, *P* < 0.001) on the scale of 4,000 km (Figure S4c), while the turnover rates were 0.077 (Figure S4c, R^2^ = 0.0158, *P* < 0.001) and 0.2191 (Figure S4c, R^2^ = 0.1185, *P* < 0.001) per 100 km in alpine and temperate grasslands, respectively. Moreover, the turnover rates for long-term environmental variables (including MAP, MAT, pH, SOC, TN and TP) were higher (0.0707, 0.2106 and 0.0181 per 100 km in the alpine grassland, temperate grassland and all sites, respectively) than those (0.0166, 0.0592 and 0.0049, per 100 km in alpine grassland, temperate grassland and all sites, respectively) for short-term environmental variables (including SWC, AP, DOC, DON, NH_4_^+^ and NO_3_^-^).

The similarity of abundant fungal taxa decreased significantly with environmental distance with the turnover rate of -0.0099 (R^2^ = 0.053, *P* < 0.001), higher than that of the entire (turnover rate = -0.0068, R^2^ = 0.053, *P* < 0.001) and rare fungal taxa (turnover rate = -0.0027, R^2^ = 0.023, *P* < 0.001) in all sites (Figure 2c). Similar environmental distance decay patterns were also found in the alpine grassland (Figure 2a), while there was no significant decay relationship for each fungal taxa in the temperate grassland (Figure 2b).

**Figure 2.**
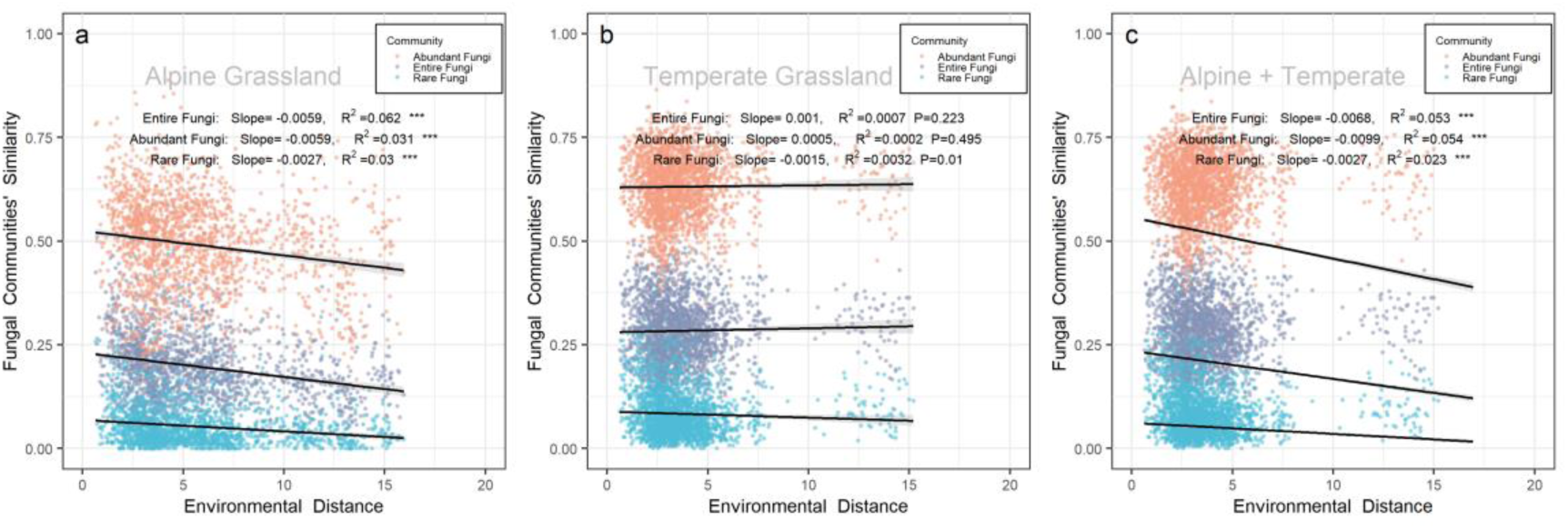
Similarities of soil entire, abundant and rare fungal taxa decreased with the increasing environmental distance in the (a) alpine grassland, (b) temperate grassland and (c) all sites. The grey, orange and blue points stand for entire, abundant and rare fungal taxa, respectively. “***” stands for significance at 0.001 level. Each fungal communities’ similarity was calculated based on binary Bray-Curtis index, and environmental distance was calculated based on Euclidean distance.

The similarity of abundant fungal taxa decreased significantly with plant communities’ dissimilarity with the turnover rate of -0.260 (R^2^ = 0.06, *P* < 0.001), which was higher than that of entire (R^2^ = 0.082, *P* < 0.001) or the rare fungal taxa (R^2^ = 0.057, *P* < 0.001) in all sites (Figure S6b). Similar distance decay patterns were also found in alpine (Figure S6c) and temperate grassland (Figure S6d) alone, though the turnover rate was higher for entire fungal taxa (turnover rate = -0.312, R^2^ = 0.190, *P* < 0.001) than that of abundant fungal taxa (turnover rate = -0.306, R^2^ = 0.090, *P* < 0.001) in the alpine grassland.

### 3.3 Soil fungal community assembly mechanisms

The null model analysis was performed to determine whether deterministic or stochastic processes dominate community assembly for entire, abundant or rare fungal taxa. The similarities of observed communities for entire fungal taxa were significantly (*P* < 0.01) higher than those of permutated communities generated by the null model in the alpine grassland, temperate grassland and all sites (Figure 3a), indicating that the entire fungal taxa were assembled by deterministic processes. The similar result was also found for abundant fungal taxa (Figure 3b), while there was no significant difference in similarities between observed and permutated communities for rare fungal taxa (Figure 3c), indicating that the rare fungal taxa was assembled by stochastic processes.

**Figure 3.**
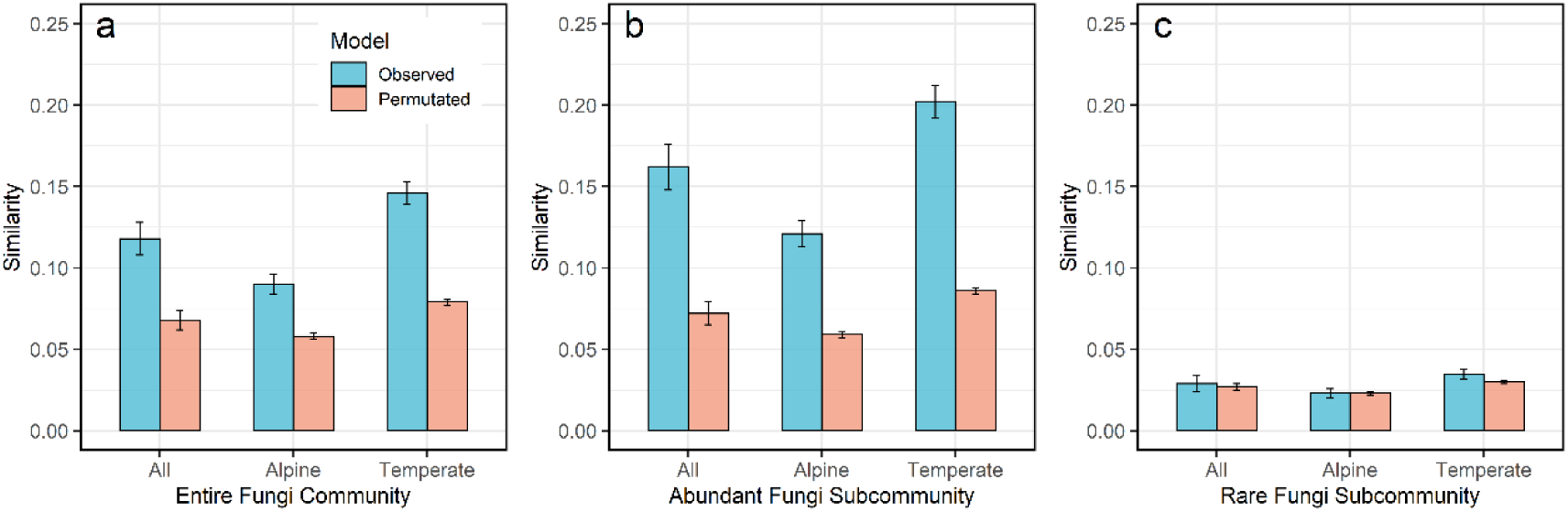
Similarity of observed (blue columns) and permutated (orange columns) soil entire (a), abundant (b) and rare (c) fungal taxa in alpine grassland, temperate grassland and all sites. The permutated similarities of entire, abundant and rare fungal taxa were analyzed by the null model analysis.

### 3.4 Deterministic factors shaping fungal community

SEM (Figure 4) showed that the similarity of the entire fungal community was directly affected by plant community similarity positively (r = 0.177), by the edaphic dissimilarity (r = -0.215), climatic dissimilarity (r = -0.048) and geographical distance (r = -0.281) negatively, and indirectly affected by geographical distance through climate dissimilarity (r = 0.068) and plant similarity (r = -0.169), respectively. Similar results were also found for abundant and rare fungal taxa with explanatory of 0.186 and 0.155, respectively. However, the impact of edaphic attributes and plant community on abundant fungal taxa (r = -0.213 and 0.166, respectively) was higher than that on rare fungal taxa (r = -0.105 and 0.059, respectively).

**Figure 4.**
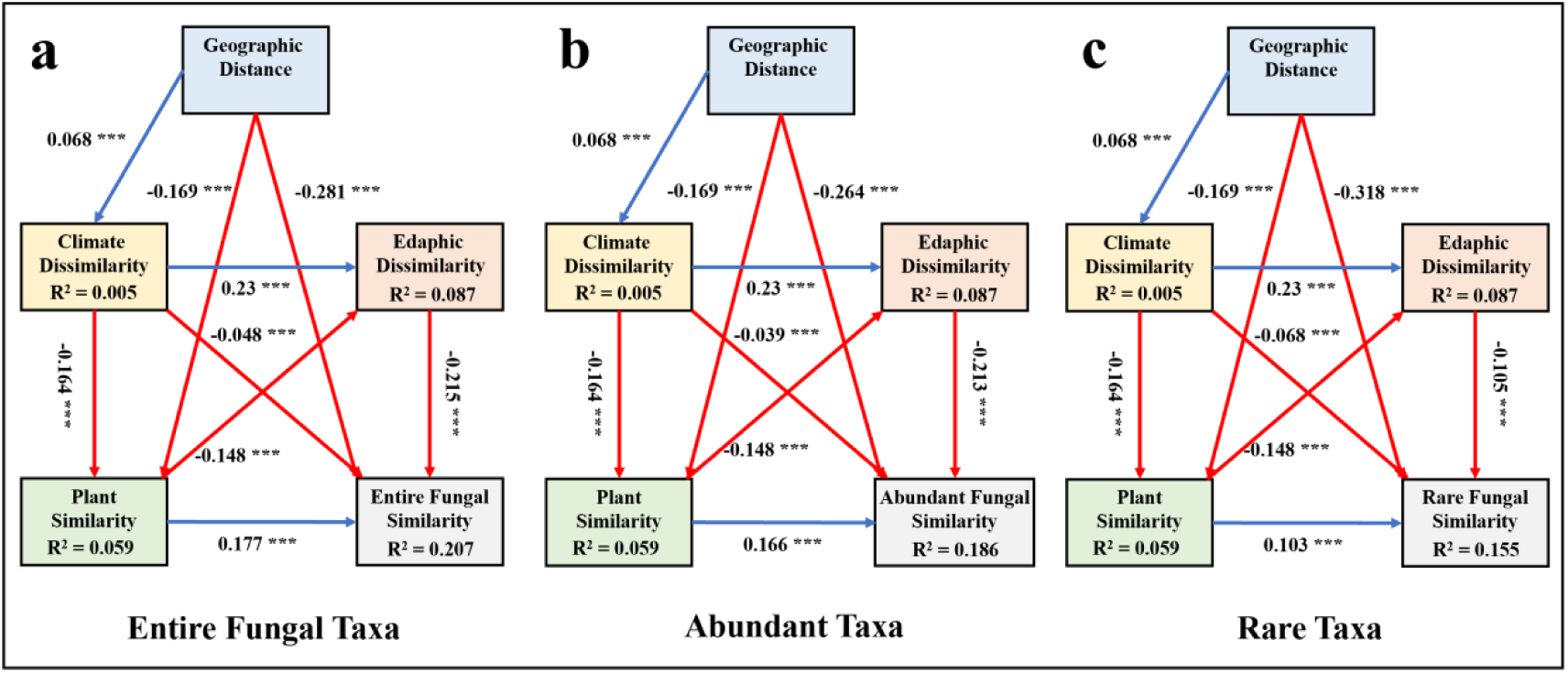
Structural equation model to quantify effects of geographic distance (latitude and longitude), climate dissimilarity (MAP and MAT), edaphic dissimilarity (SOC, DOC, TN, TON, NH_4_^+^, NO_3_^-^, TP, AP, SWC and pH), and plant community dissimilarity on soil (a) entire, (b) abundant and (c) rare taxa in all sites. Red and blue lines stand for negative and positive correlations, respectively. The solid arrow represents significant correlations (the number is the r values; the asterisk indicates the p-value: *, *P* < 0.05; **, *P* < 0.01; ***, *P* < 0.001).

### 3.5 Dispersal potential and niche breadth of entire, abundant and rare fungal taxa

The immigration rates of fungal taxa in the temperate grassland (Figure 5a, 0.042 ± 0.015, 0.739 ± 0.039 and 0.111 ± 0.005 for entire, abundant and rare fungal taxa, respectively) were significantly higher than those in the alpine grassland (0.025 ± 0.011, 0.525 ± 0.045 and 0.088 ± 0.005 for entire, abundant and rare fungal taxa, respectively). Moreover, the dispersal rate of abundant fungal taxa was far higher than that of rare fungal taxa either in the alpine, temperate grassland or all sites.

**Figure 5.**
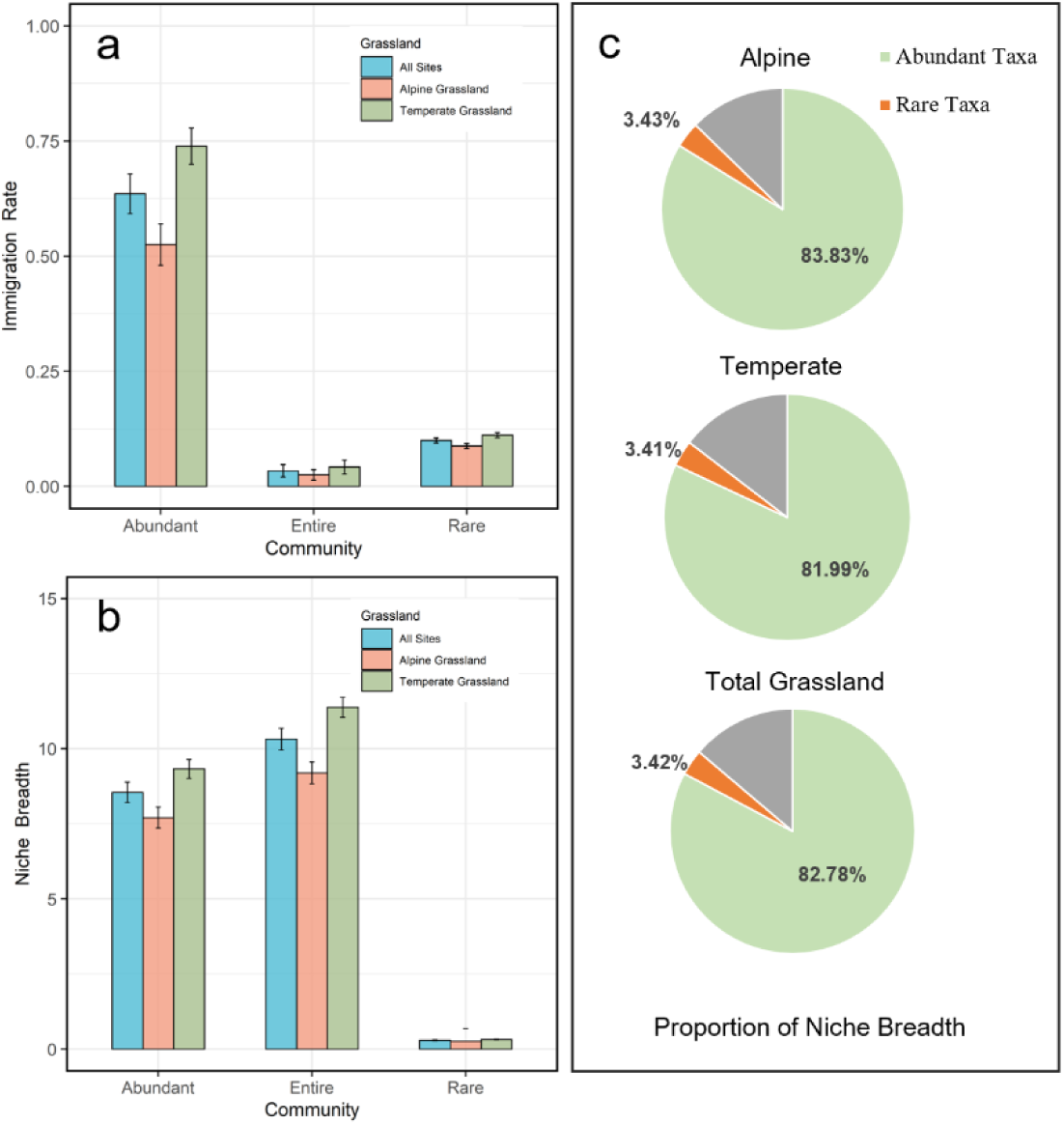
The immigration rate (a), niche breadth (b) and the proportion of niche breadth (c) of entire, abundant and rare fungal taxa in the alpine grassland, temperate grassland, and all sites. Immigration rates were calculated by Etienne’s method; the ecological niche breadth was calculated by Levins’ method. The proportions of niche breadth for abundant and rare fungal taxa were calculated by their ratios to the niche breadth of the entire fungal taxa.

The niche breadth of fungal taxa in the temperate grassland (11.38 ± 0.33, 9.33 ± 0.31 and 0.32 ± 0.01 for entire, abundant and rare fungal taxa, respectively) was significantly higher than that in the alpine grassland (9.20 ± 0.36, 7.71 ± 0.35 and 0.26 ± 0.41 for entire, abundant and rare fungal taxa, respectively), and the contribution of abundant taxa to the community niche breadth was as high as 83.83 % (Figure 4 b and c). In contrast, the contributions of the rare fungal taxa were only 3.43%, 3.41% and 3.42% in alpine grassland, temperate grassland and all sites, though rare OTUs (5, 779) accounting for 63.02% of the total OTUs.

## Discussion

In 129 soil samples collected across 4,000 km in Chinese Northern grasslands, we found that both rare and abundant fungal taxa decayed with the increasing geographic distance, though the rare taxa had a much lower spatial turnover rate. The spatial pattern and turnover of microbial community is one of the fundamental topics in biogeography and microbial ecology, linked with biodiversity shaping and maintaining (Hanson et al. 2012, Bahram et al. 2013). Previously, we found that the soil entire prokaryotic taxa exhibited a U shape pattern with geographic distance across alpine and temperate grasslands (Zhang et al. 2022). In this study, the entire soil fungal taxa exhibited a distance-decay pattern for the same set of samples. The distance-decay pattern of soil prokaryotes was mainly influenced by short-term environmental variables such as soil water and dissolved nutrients (Zhang et al. 2022), which existed a U shape pattern with geographic distance. In contrast, fungi are more resistant to water disturbance than bacteria since the fungal hyphae could help fungi to capture more water and dissolved nutrients from disturbed environment, thus making them be less responsive to their changes (Treseder and Lennonb 2015, Frac et al. 2018, Crowther et al. 2019). Thus, unlike prokaryotes, the fungal community was less influenced or at least not dominantly influenced by short-term environmental variables with a core of soil water, and exhibited a decay relationship with geographic distance as well as some other previous studies (Tedersoo et al. 2014, Ma et al. 2017).

However, the turnover rates for soil fungi (Figure S9, -0.003 and -0.006 per 100 km in alpine and temperate grasslands, respectively) are lower than those of the corresponding prokaryotic communities (-0.005 and -0.012 per 100 km in alpine and temperate grasslands, respectively). The lower turnover rates for fungi is consistent with the study in the boreal forest soil (Ma et al. 2017). The lower turnover rates of soil fungi than prokaryote could be attributed to differences in their body sizes, growth rates, and dispersal capabilities (Tedersoo et al. 2014, Peay et al. 2016). Moreover, fungi could alleviate the stresses such as water limitation and nutrient limitation on their distribution, leading to a relatively low turnover rate (Tedersoo et al. 2014, Frac et al. 2018).

Different from the entire and abundant fungal taxa that were assembled by deterministic processes, rare fungal taxa were assembled mainly by stochastic mechanisms. Stochastic processes were less influenced by environmental variance along the geographic distance (Zhou and Ning 2017); while deterministic factors including soil variables, climate and plant traits changed greatly along the geographic distance (Figure S4, S5 and S6). Consistently, the abundant fungal taxa were reported to be more sensitive to environmental changes spatially since their turnover rate by the environmental distance was much higher than that of rare taxa (Wang et al. 2022a, Wang et al. 2023a). Dominant species may have higher environmental adaptability (Jousset et al. 2017, Jia et al. 2018) to these changes in environmental factors along the geographic distance. Consistently, as proved by the niche breadth analysis, the niche breadth of abundant taxa was far higher than that of rare taxa. In contrast, rare species are likely with weak environmental adaptability, thus was largely shaped by ecological drift, like the random appearance of rare species likely due to their high richness but low occupancy and abundance (Jia et al. 2018, Pascoal et al. 2021, Ren and Gao 2022). Inconsistently, some studies found that rare taxa were dominated by deterministic processes in an agriculture ecosystem (Jiao and Lu 2020a) or an alpine grassland ecosystem at the scale of 2,000 km (Tang et al. 2023), likely due to different ecosystem or scales were investigated.

Moreover, the lower turnover rate of rare fungal taxa than that of abundant fungal taxa was observed in this study. Opposite to a higher turnover rate for rare than abundant fungal taxa in previous studies (Jiao and Lu 2020a, Wang et al. 2022a, Tang et al. 2023, Wang et al. 2023a). The lower turnover may be caused by the difference of their responses to environments change, tolerance ability (Jia et al. 2018, Jiao and Lu 2020b), and dispersal potential (Jiao and Lu 2020a, Wang et al. 2022b). Firstly, abundant fungal taxa had higher immigration rates than rare fungal taxa either in the alpine or temperate grassland (Figure 5a) in this study. Secondly, the higher frequency of abundant OTUs on average than rare OTUs proved the difference in dispersal potential between abundant and rare taxa. Rarity could be associated with taxa occurring at narrow environmental ranges or dispersal limitation (Sul et al. 2013, Jia et al. 2018). With low dispersal potential, the dispersal limitation combined with drift could lead to a large community turnover for rare fungal taxa (Leibold et al. 2004, Jia et al. 2018). While the high dispersal potential of abundant fungal taxa implies more species exchange and leads to high similarity among sites. Moreover, rare fungal taxa were less sensitive to environmental change compared to abundant taxa as shown in this and other studies (Jiao and Lu 2020a, Wan et al. 2021), thus contributing to lower turnover rates as well.

## Conclusion

Our study revealed soil fungal spatial patterns and underlying mechanisms in grassland ecosystems for entire, abundant, and rare fungal taxa on the scale of 4000 km in Chinese Northern Grasslands. Unlike the U pattern for soil prokaryotes, soil entire fungal taxa exhibited significant distance-decay patterns, and so did abundant and rare fungal taxa. Moreover, the turnover rate for rare taxa was lower than abundant fungal taxa, which was attributed to their distinct assembly mechanisms. The rare fungal taxa were assembled mainly by stochastic processes, that were less influenced along geographic distance. While abundant fungal taxa were assembled mainly by deterministic processes, which changed greatly along the geographic distance. In summary, our results provide insights into the distinct spatial pattern between rare and abundant fungal taxa at a large scale in Chinese Northern grasslands.

## Acknowledgement

This work was financially supported by the Second Tibetan Plateau Scientific Expedition and Research (STEP) Program (Grant No. 2019QZKK0304-02), Joint Chinese Academy of Sciences (CAS)-Max Planck Gesellschaft (MPG) Research Project (HZXM20225001MI), the Scientific research ability improvement project for outstanding teachers of Chinese Academy of Sciences (E1E40513X2), the Joint Research on Ecological Conservation and High- Quality Development of the Yellow River Basin program (2022-YRUC-01-0102), National Natural Science Foundation of China (42041005), CAS “Light of West China” Program, and the Fundamental Research Funds for the Central Universities.

## Data availability statement

The soil fungi dataset has been deposited in the NCBI Sequence Read Archive under accession number: PRJNA1129116.

## Conflict of interest statement

The authors declare that they have no known competing financial interests or personal relationships that could have appeared to influence the work reported in this paper.

